# Effects of Blood Sugar Levels, Kidney Disease and Medication on Anaemia Status of Type 2 Diabetes Mellitus Patients

**DOI:** 10.1101/2020.03.21.001172

**Authors:** Malala Mulavu, Mildred Zulu, Musalula Sinkala, Sody Munsaka, Pauline Okuku, Panji Nkhoma

## Abstract

**Introduction:** Lifestyle-related diseases like type 2 diabetes mellitus, have emerged as a significant public health problem due to rapid urbanization and industrialization. In 2018, the prevalence of type 2 diabetes mellitus was estimated to be 500 million cases worldwide and was comparable between high- and low-income countries. Diabetes leads to multiple complications, including end-stage renal disease, cardiovascular disease, infection, and death. Anaemia in diabetic individuals has severe adverse effects on the quality of life and is associated with disease progression. A cross-sectional study was conducted that included 101 participants. The study aimed to determine the prevalence of anaemia and kidney disease and their interplay with medication and/or blood glucose levels in T2D patients.

**Results:** seventy-one per cent of participants were females. Majority of patients 62% were on Insulin, 30% on metformin, 7% on a combination of metformin and glimepiride and 1% on glimepiride. Ninety-five (94%) of the participants were HIV negative. The prevalence of anaemia among the participants was 23% out of which 56% had moderate anaemia, and 44% had mild anaemia. Twenty-one per cent (21%) of the participants had high creatinine levels signifying impaired kidney function or kidney disease. Anaemia was significantly associated with kidney function, fasting blood glucose and use of metformin; p = 0.042 beta = 2.5, p = 0.025 beta = 2.7 and p = 0.040 beta = −2.5 respectively.

**Conclusion:** The study found the prevalence of anaemia of 23%, which was of moderate public health concern. Also, the prevalence of kidney disease was high in patients with Type 2 Diabetes Mellitus. It also found that kidney disease and high blood glucose levels increase the chances of developing anaemia. However, we found that metformin had a protective role against the development of anaemia in Type 2 Diabetes Mellitus patients.

## 1.0 INTRODUCTION

Lifestyle-related diseases like diabetes mellitus, have emerged as a significant public health problem due to rapid urbanization and industrialization [1]. In 2018, the prevalence of type 2 diabetes mellitus (T2D) was estimated to be 500 million cases worldwide and was comparable between high and low-income countries [2]. However, others have suggested that the condition may be up to two times more prevalent in low-income populations compared to wealthy populations [3–5]. The number of adults with diabetes mellitus in sub-Saharan Africa is predicted to rise from 19.8 million in 2013 to 41.5 million in 2035 [6–10], and in 2016, the age-standardised prevalence of diabetes in Zambia was 3.5% [11].

Diabetes is the leading causes of kidney disease [12]. About 1 out of 4 adults with diabetes have kidney disease [13]. It leads to multiple complications, including end-stage renal disease, cardiovascular disease, infection, and death [14]. In the recent past, the prevalence of diabetes and diabetic kidney disease has increased [15].

Anaemia is a reduction in the haemoglobin concentration of the blood below normal for age and sex, and the World Health Organisation defines anaemia as Hb level < 13 g/dL in men and 12 g/dL in women [16]. Anaemia in diabetic individuals has severe adverse effects on the quality of life and is associated with disease progression and the development of comorbidities [17]. Obesity and dyslipidaemias, which are strongly associated with the diabetic framework, significantly contribute to increasing the risk of cardiovascular diseases [18]. The study aimed to determine the prevalence of anaemia and kidney disease and their interplay with medication and/or blood glucose levels in T2D patients at the University Teaching Hospital, which is the largest third-level referral in Zambia.

## 2.0 MATERIALS AND METHODS

### 2.1 Study design

The study was cross-sectional and was carried out at the University Teaching Hospital, a tertiary hospital in Lusaka, Zambia.

### 2.2 Study Population

The study population included all existing and newly diagnosed type 2 diabetes mellitus patients visiting UTH from April to July 2019. We used convenience sampling to select participants.

### 2.3 Specimen Collection

Blood samples were collected by phlebotomy. From each patient 3ml of blood, were collected in each of two tubes; one Ethylenediaminetetraacetic acid (EDTA) for haematological analysis, and lithium heparin tube for biochemical analysis (kidney function tests).

### 2.4 Full Blood Count, Kidney Function and Fasting Blood Sugar

Full blood count was measured using Sysmex XT-4000i (Germany) automated analyser. Kidney function measurements were carried out using the ABX Pentra 400, Japanese chemistry analyzer. Glucose was measured using an Accu check™ glucometer at the point of specimen collection. The patient’s clinical records were collected using the UTH laboratory information system. All quality control procedures were followed before running the specimens.

### 2.5 Statistical Analysis

The data were entered in Excel & analysed using Python 3.7 for mac. Continuous variables were summarised using mean, standard deviation, percentiles, range and mode. Descriptive statistics in the form of percentages and graphs were used to show the prevalence of anaemia and kidney disease in type 2 diabetic mellitus patients. Multiple logistic regression was used to determine the effects of Kidney function, fasting blood glucose and medication on the occurrence of anaemia and the Student t-test was used to determine the mean differences in the kidney function tests between study participants and the non-diabetic control group.

### 2.6 Ethical Approval

**Ethical approval for the study was obtained from The University of Zambia Health Science Research Ethics Committee (UNZAHSREC) (Protocol ID: 20190117001 of IORG0009227). Informed Consent was obtained from all study participants.**

## 3.0 RESULTS

### 3.1 Demographic Characteristics

One hundred and one participants took part in the study out of which 71 (70%) were females. Minimum age of participants was 22 years, maximum of 83 years with a mean age of 54.43 ± 14 years. The majority of patients 63 (62%) were on Insulin, 30 (30%) on metformin, 7 (7%) on a combination of metformin and glimepiride and only 1 (1%) on glimepiride (Figure 1). Ninety-five (94%) of the participants were HIV negative.

**Figure 1:**
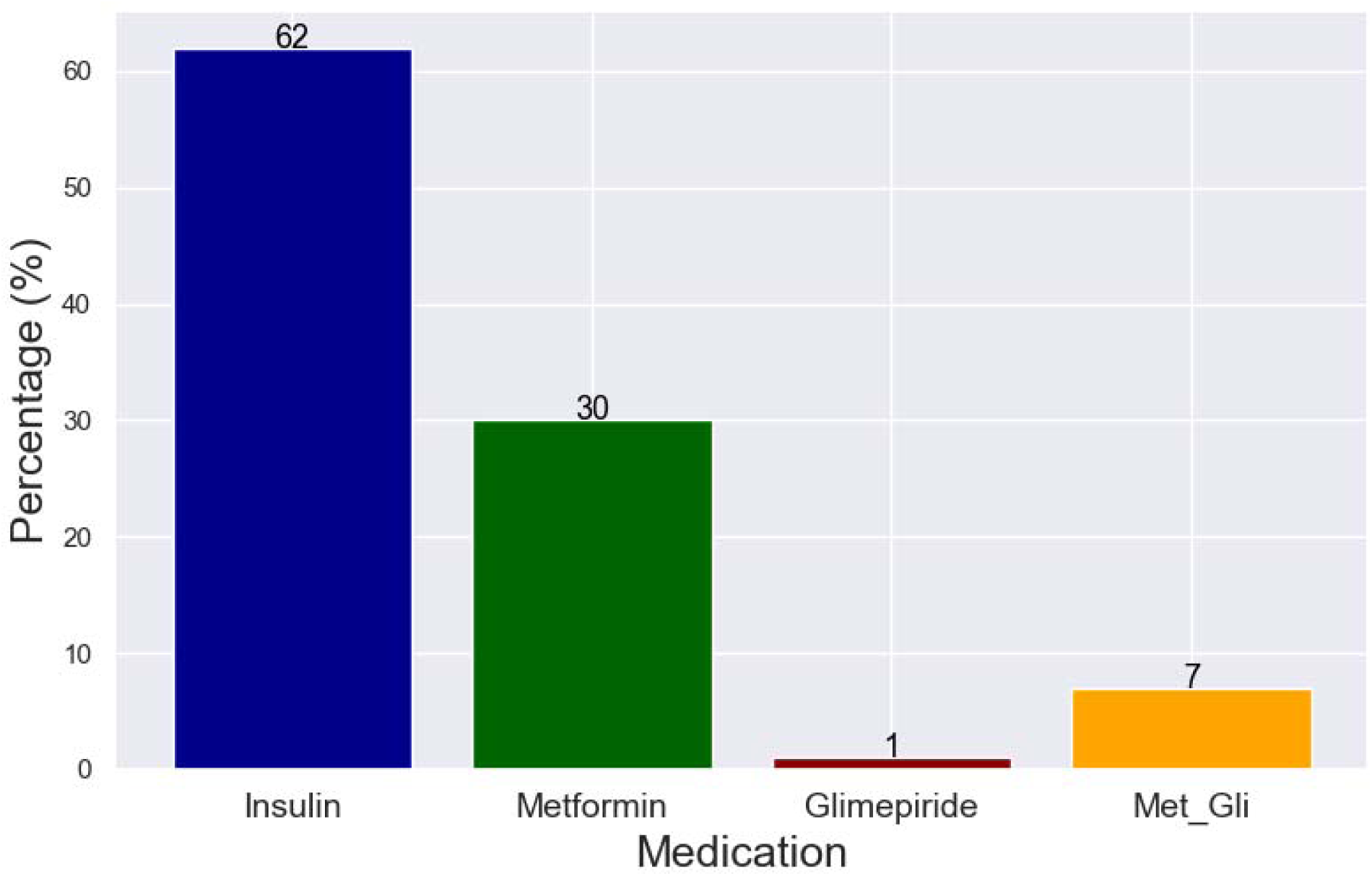
Distribution of Antidiabetic medication of participants.

### 3.2 Fasting Blood Sugar

The minimum fasting blood glucose recorded was 3.8mmol/l, maximum 24.9 mmol/l with a mean blood glucose of 10.24 ± 4.42mmol/l. 78% had higher than normal fasting blood glucose levels. 22% had normal blood glucose levels, and none had low blood glucose levels (Figure 2).

**Figure 2:**
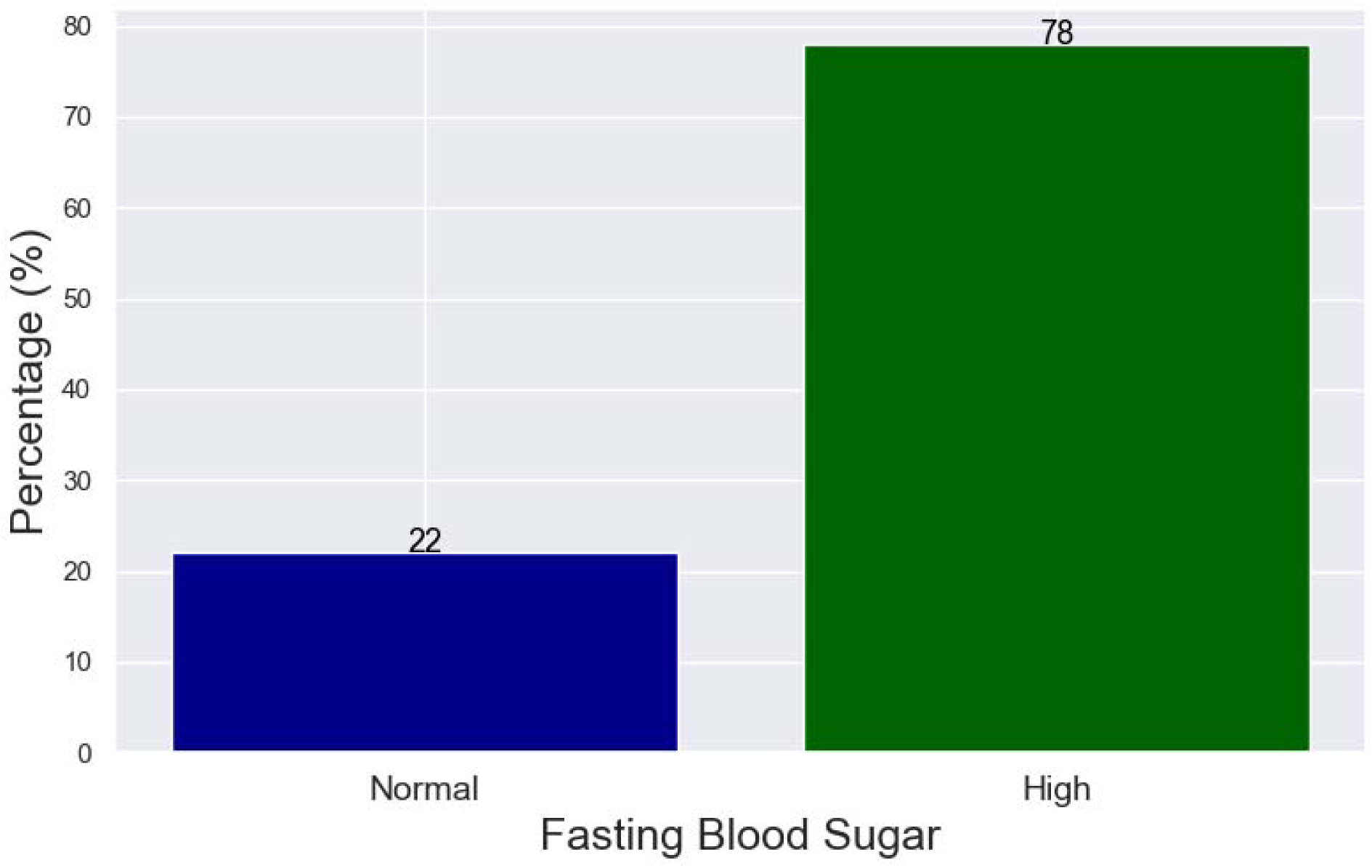
Fasting blood glucose levels of participants.

### 3.3 Anaemia

The prevalence of anaemia among the participants was 23%, of which 23 out of the 101 participants were anaemic (figure 3). The minimum haemoglobin concentration was 9.7g/dl, and the maximum was 17.2g/dl with the mean haemoglobin concentration of 13.75 ± 1.63 g/dl. Concerning the severity of anaemia, we found that 56% of the anaemic patients had moderate anaemia, 44% had mild anaemia with none having severe anaemia (Figure 4). We defined the severity and classification of anaemia based on the World Health Organization (WHO) criteria [16] Out of the 23 anaemic participants, 67% had microcytic, hypochromic anaemia, 33 % had normochromic normocytic anaemia, and none of them had macrocytic anaemia.

**Figure 3:**
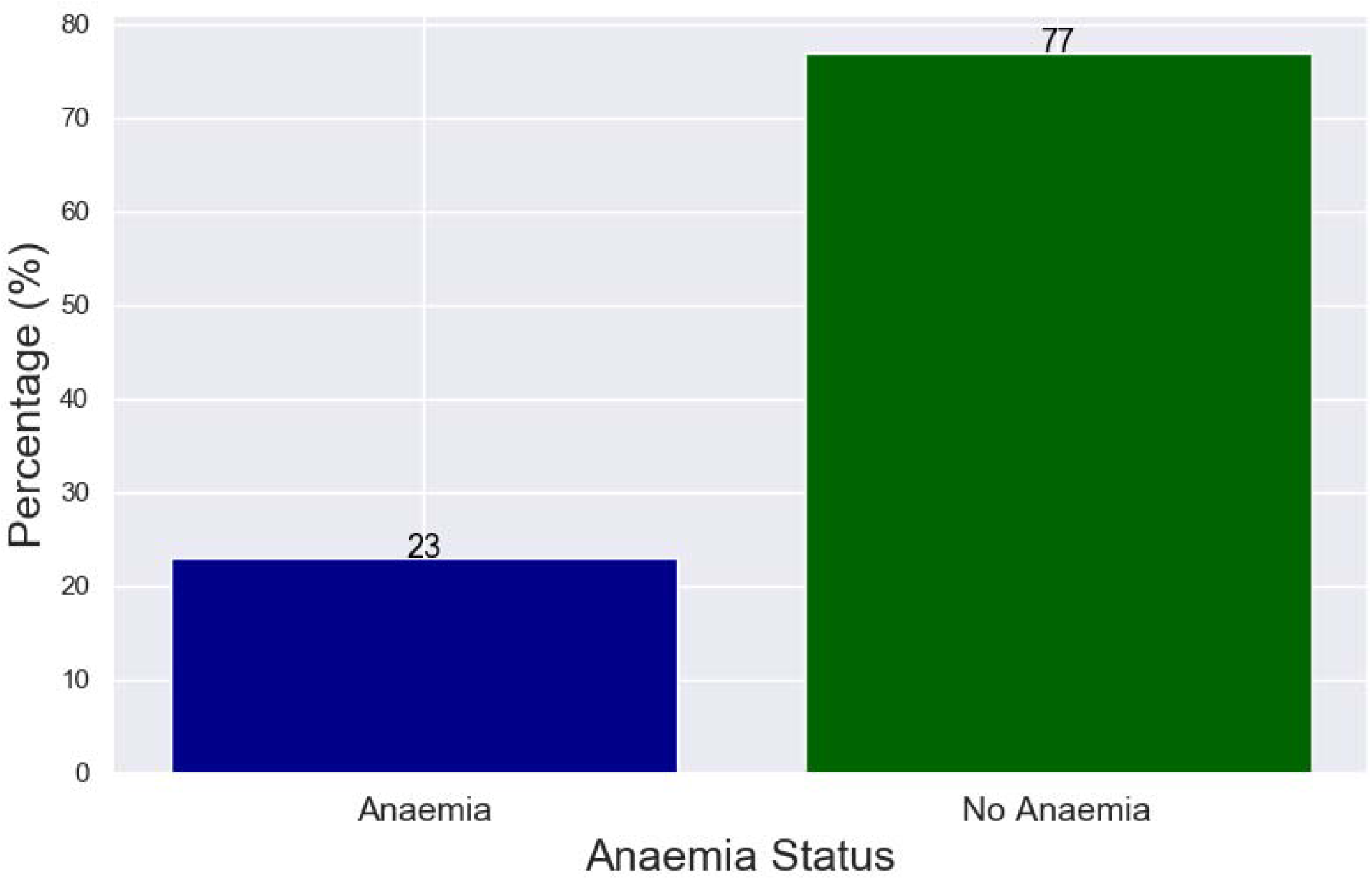
Prevalence of anaemia.

**Figure 4:**
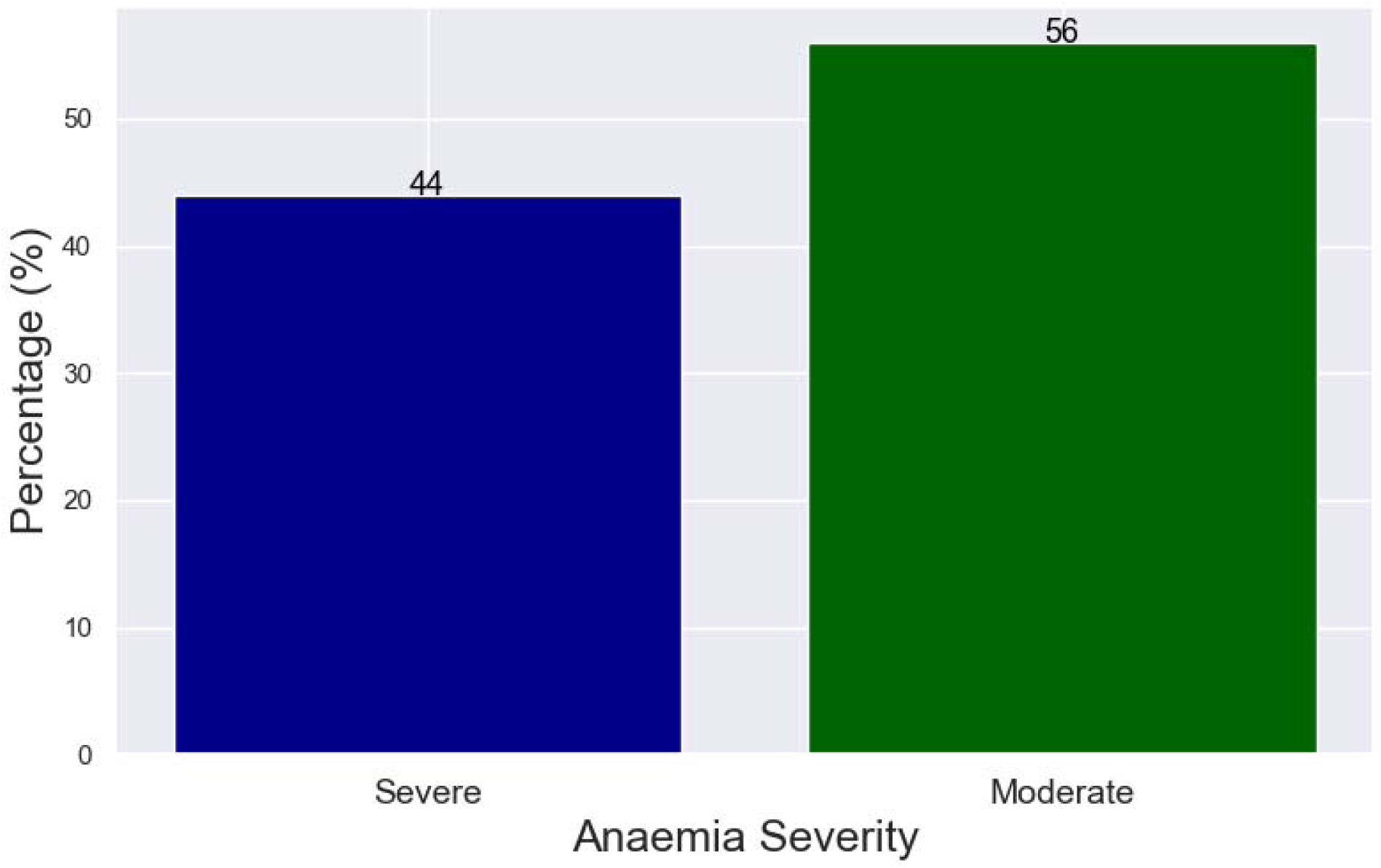
Anaemia severity.

### 3.4 Kidney Function

We found that twenty-one per cent (21%) of the participants had higher than normal (110μmol/l) creatinine levels signifying impaired kidney function or kidney disease (Figure 5). The minimum creatinine level that we recorded was 50.8 μmol/l, while the maximum level was at 404.8 μmol/l with a mean of 100.77 ± 55.96 μmol/l. We used creatine to assess kidney function because serum creatinine above specific cutoff points reliably identifies patients with acute kidney injury or chronic kidney disease [19,20]. Anaemia was significantly associated with kidney function (p = 0.042; beta = 2.5), fasting blood glucose (p = 0.025; beta = 2.7) and use of metformin (p = 0.040; beta = - 2.5)

**Figure 5:**
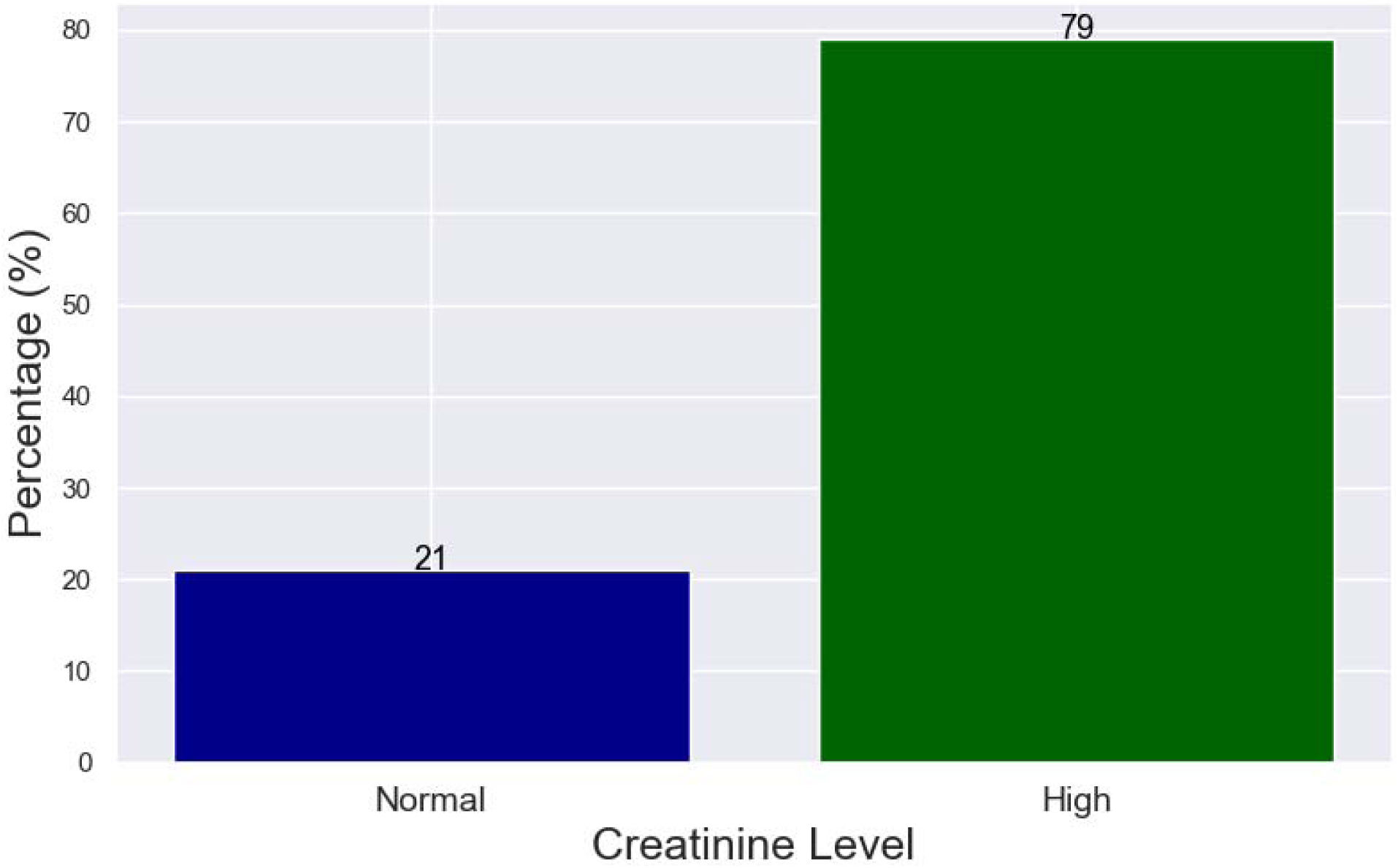
Prevalence of kidney disease.

After Adjusting for the potential confounding factors; sex, alcohol consumption and HIV status:

i. We found that the odds of having anaemia were 11.76 times higher for participants with kidney malfunction (OR = 11.76, 95% CI =1.09 - 126.68, p = 0.042).
ii. Furthermore, we showed that the odds of having anaemia were 15.45 times higher for participants with high fasting blood glucose (OR = 15.45, 95% CI =1.40 - 170.37, p = 0.025).
iii. For the participants taking metformin, we found that their odds of having anaemia were 0.09 times less, and showed a negative beta value of −2.5 (OR = 0.09, 95% CI = 0.01 - 0.89, p = 0.040), denoting that these participants on metformin were about 91% protected from developing anaemia.

We compared the plasma creatinine, urea, sodium, chloride and potassium levels between the diabetic participants to those of the non-diabetic individuals. Here, we found that the creatinine (p = 1.36 x 10^−6^), urea (p = 0.011), sodium (p = 1.84 x 10^−9^) and chloride (p = 2.59 x 10^−7^) levels were significantly higher in the diabetic study participants than in the non-diabetic control group (Figure 6 and 7). However, potassium ion levels between the two groups were not statistically significant, with a p-value of 0.8222.

**Figure 6:**
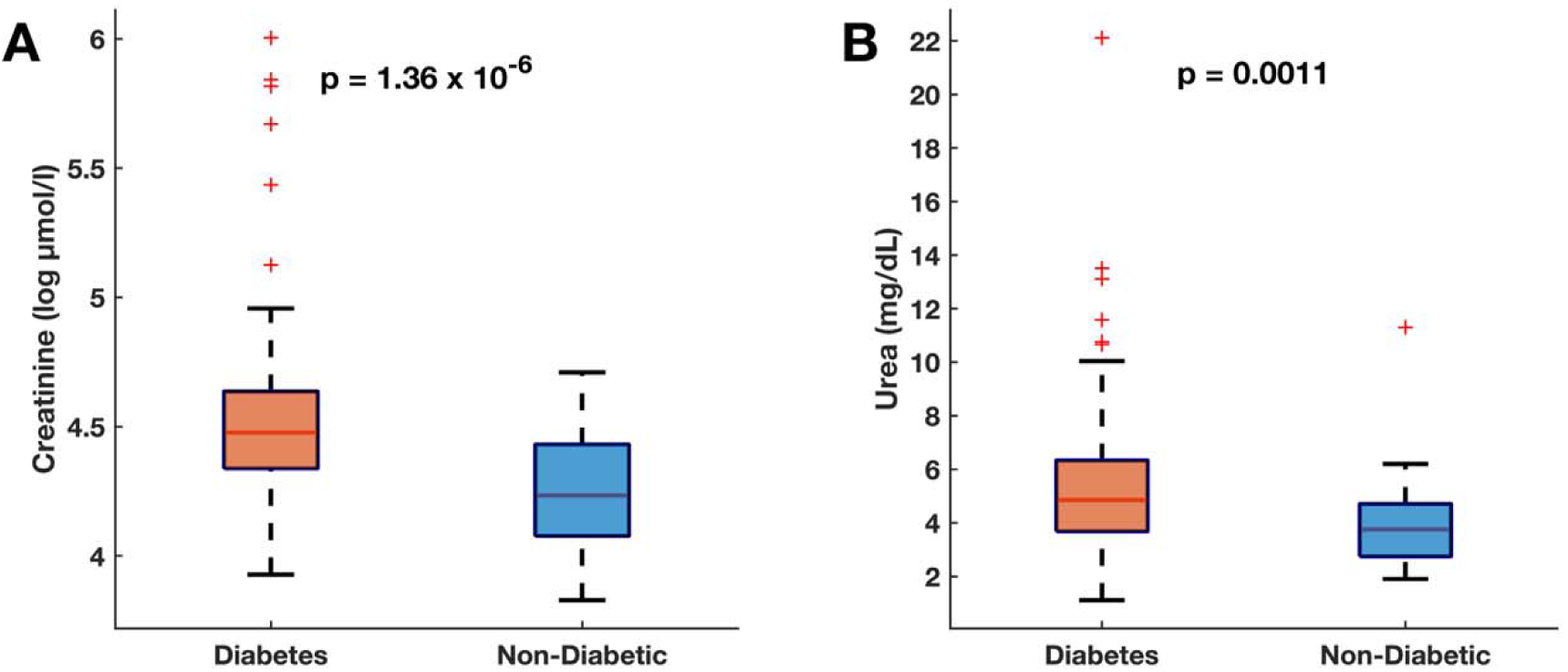
Mean Differences in creatinine and urea between diabetic participants and non-diabetic controls.

**Figure 7:**
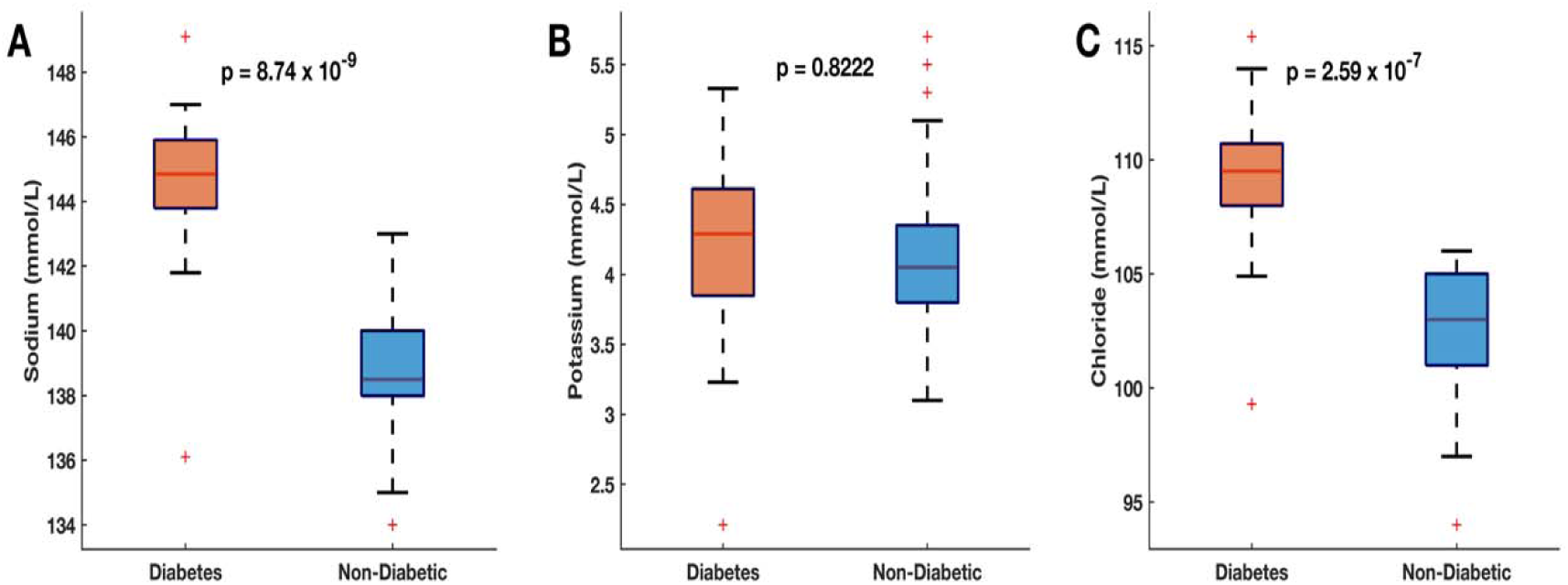
Mean Differences in sodium, potassium and chloride ions between diabetic participants and non-diabetic controls.

## 4.0 DISCUSSION

Diabetes Mellitus is frequently accompanied by mild to moderate anaemia, which is often called anaemia of inflammation or infection or anaemia of chronic disease [21]. The reduced Hb levels identify diabetic patients at increased risk for hospitalization and premature death [22]. Our study found that nearly one in four diabetic patients have anaemia which, according to WHO is of moderate public health significance [23]. Others have also found comparable prevalences of anaemia in India (18%) and Kuwait (29.7%) [1,24].

The predominant risk factor for the development of anaemia in a diabetic population is the presence of renal disease, which manifests as either impaired renal function or albuminuria [25]. In particular, the critical factor underlying the development of anaemia is the uncoupling of the haemoglobin concentration from renal erythropoietin synthesis [24]. In our study, t[26][25]he high incidence of anaemia may also be due to other risk factors that are related to DM. Several studies have reported factors that increase the risk of anaemia, which include; damage to renal interstitium due to chronic hyperglycaemia and consequent formation of advanced glycation end products by increased reactive oxygen species, and systemic inflammation as well as reduced androgen levels induced by diabetes [27]. Here also, the increased levels of inflammatory cytokines (IL 6) may promote the apoptosis of immature erythrocytes, thus reducing the number of circulating erythrocytes and haemoglobin levels [17,28]. We revealed that the majority of participants had microcytic hypochromic anaemia. Our findings differ from those obtained in a study done in Ghana, where they found that; the majority of participants had normochromic normocytic anaemia, followed by hypochromic microcytic, and lastly, normochromic macrocytic anaemia [29]. Here, the observed difference between the findings Ghana study and our study may be due to the differences in the duration of illness. In our study, the majority of the participants had diabetes for less than five years. Conversely, the part participants in the Ghana study may have had diabetes for a more extended period resulting in normochromic normocytic anaemia which is commonly associated with anaemia of chronic diseases such as diabetes.

We found that none of our study participants had severe anaemia, but the majority had moderate anaemia. We suggest that since our study participants were diagnosed with diabetes within the previous five years, it is unlikely that they would develop increased levels of inflammation with decreased erythropoietin levels both which cause severe anaemia in diabetes. Here, others have also shown that people with diabetes have mild to moderate anaemia [1,30].

Kidney disease is a common occurrence in patients with T2D. Approximately 30% of patients with diabetic nephropathy eventually progress to end-stage renal failure, and the rest usually die from cardiovascular disease before reaching end-stage [31]. The increased blood glucose level that leads to damage of the kidney glomerular capsules is a significant cause of kidney disease in diabetics. Here also, high blood pressure can damage the kidney blood vessels, thereby leading to glomerular failure.

Our results revealed that 21% of participants had an elevated serum creatinine. Here, our finding is similar to those of a study done in the USA, where they found a prevalence of CKD of 24.1% in T2D patients [32]. Another similar study carried out in Finland showed that 68.9% of patients presented with signs of CKD out of which 34.7% appeared to suffer from significant CKD [33]. Another study done in Zimbabwe on African diabetics found that 44.8% of the patients suffered from nephropathy [34]. All the highlighted studies demonstrated a much higher prevalence of CKD compared to our study. Here also, the majority of our study participants were diagnosed with diabetes within five years; therefore, had little damage to the glomerulus and did not have many complications that are associated with high blood pressure. Creatinine, urea and electrolytes (sodium and chloride) were higher in the diabetic group compared to the non-diabetic control group. These findings are consistent with the current literature, which indicates that electrolyte imbalance is a common feature of diabetes mellitus associated with hyperglycaemia [35–37]. Furthermore, the observed lower levels of potassium in the diabetic group compared to the non-diabetic group can be explained by the noted high levels of insulin and glucose in the diabetic group, both of which promote the cellular influx of potassium from extracellular compartment [38–40].

We found that participants with high fasting blood glucose were significantly more likely to develop anaemia. We suggest that this may be because chronic hyperglycaemia may result in abnormal red blood cells, oxidative stress, and sympathetic denervation of the kidney related to autonomic neuropathy. These factors promote a hypoxic environment in the renal interstitium, which leads to impaired production of erythropoietin by the peritubular fibroblasts [27]. This suggests that the incidence of anaemia is likely to increase in poorly controlled diabetes, and therefore reducing blood glucose levels could assist in the reduction of the risk of anaemia in diabetic populations.

We also found that the odds of having anaemia were 0.09 times less for participants taking metformin. The odds ratio for metformin is less than one, which means that metformin significantly reduces the chances of developing anaemia in diabetes individuals and therefore plays a protective role. This is contrary to studies that have suggested that metformin leads to anaemia in patients with diabetes due to the resulting vitamin B12 deficiency that is caused by prolonged use of the drug [41]. This difference may be due to vitamin B12 deficiency anaemia that occurs after prolonged use, while the majority of our study participants had diabetes for less than 5yrs. Therefore, they may not have used metformin for an extended period.

## 5.0 CONCLUSION

We found that there is a prevalence of 23% of anaemia and 21% of kidney disease in patients with type 2 diabetes mellitus at the University Teaching Hospital. Furthermore, we found that patients with kidney disease and increased blood glucose levels have an increased chance of developing anaemia. Also, we found that metformin played a protective role in the development of anaemia in our study participants. We suggest, therefore, that among T2D patients, it is essential to monitor regularly for anaemia and kidney disease the two conditions are typical ofT2D patients.

## 8.0 AUTHOR CONTRIBUTIONS

Conceptualisation: Panji Nkhoma, Mildred Zulu, Musalula SInkala, Malala Mulavu.

Data Analysis: Panji Nkhoma, Musalula Sinkala, Sody Munsaka.

Investigation: Malala Mulavu, Pauline Okuku

Supervision: Panji Nkhoma, Mildred Zulu, Musalula Sinkala, Sody Munsaka

Writing original draft: Panji Nkhoma, Mildred Zulu, Sody Munsaka, Musalula Sinkala

